## Supplementary figures and images for "Negative cell cycle regulation by Calcineurin is necessary for proper beta cell regeneration in zebrafish"

### Supplemental figure 1

#### Figure 1 supplemental

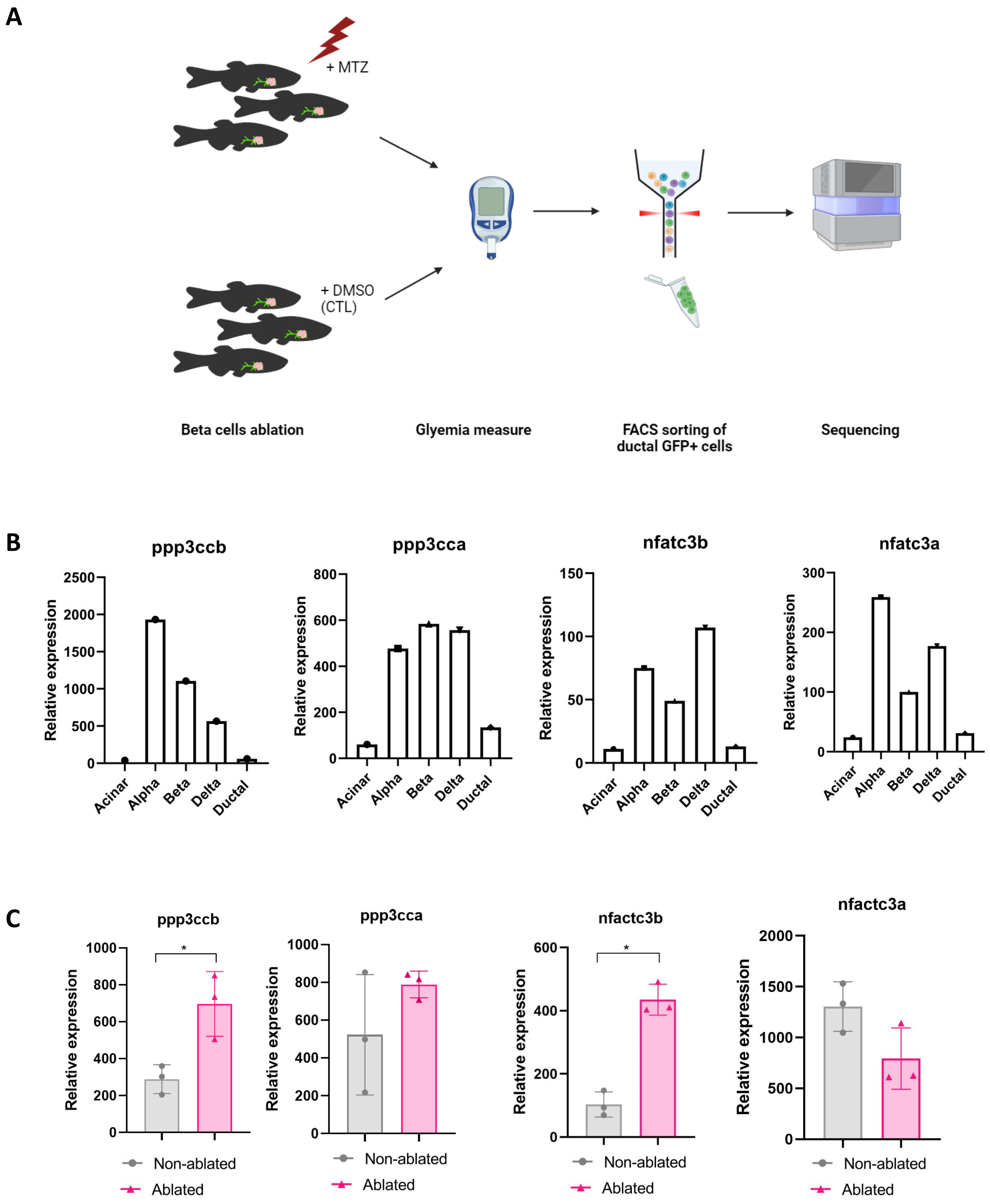

### Supplemental figure 2

#### Figure 2 supplemental

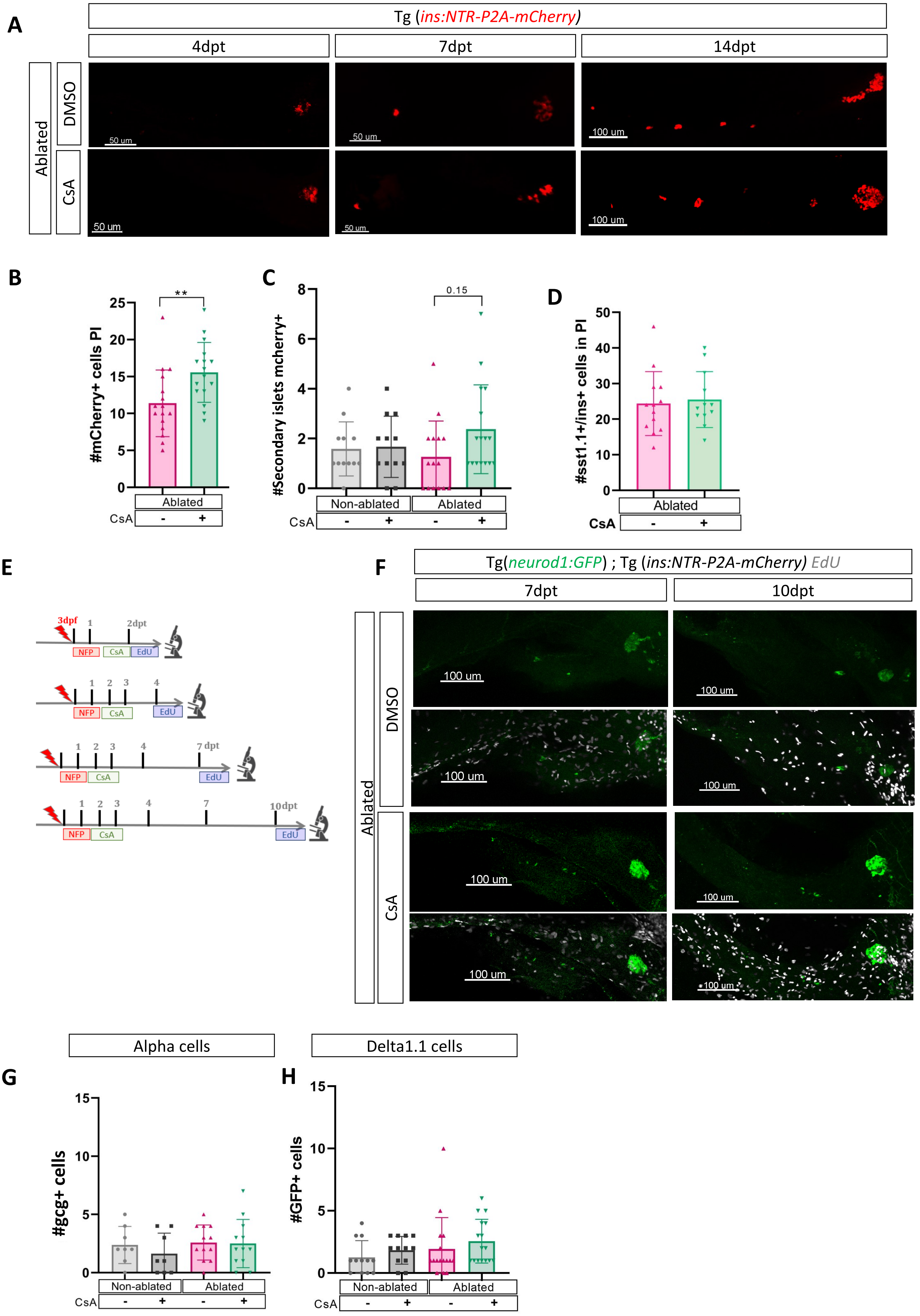

### Supplemental figure 4

#### Figure 4 supplemental

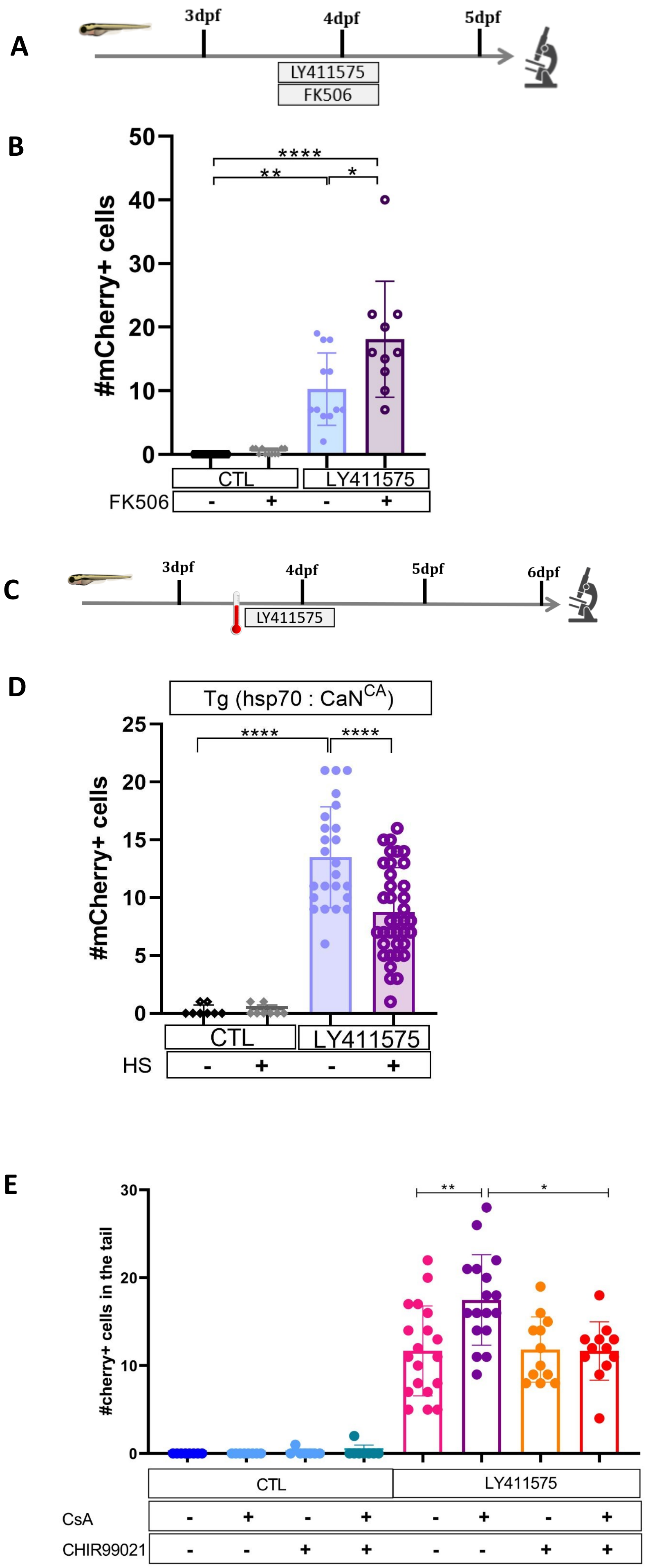

### Supplemental figure 5

#### Figure 5 supplemental

A

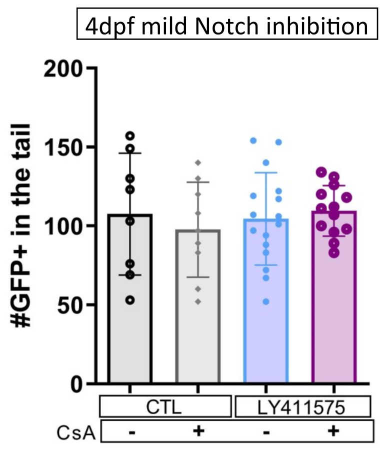


B

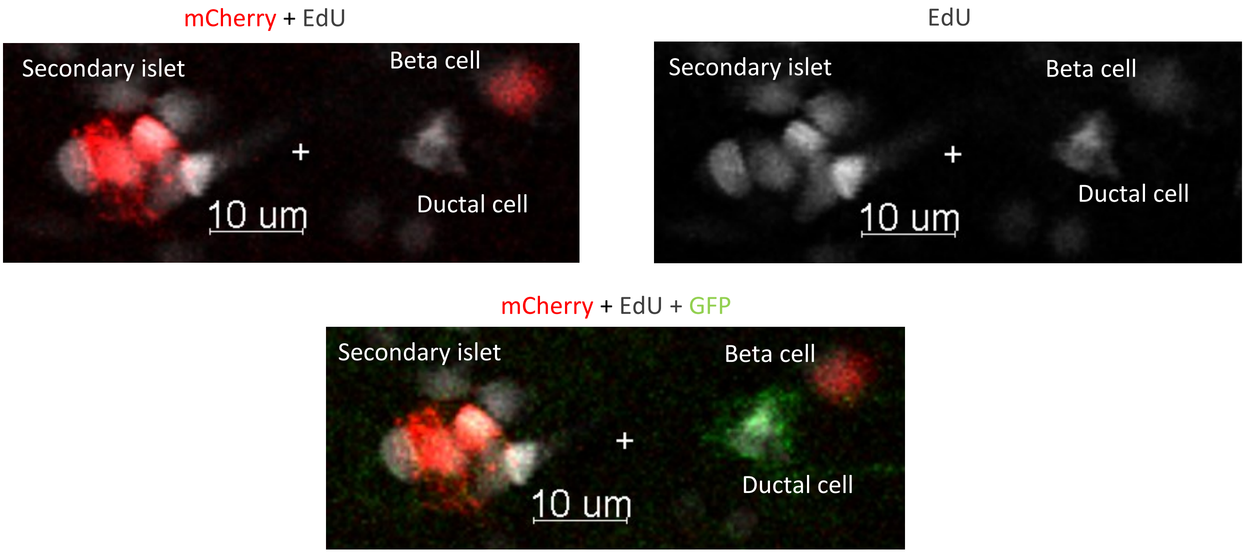


C

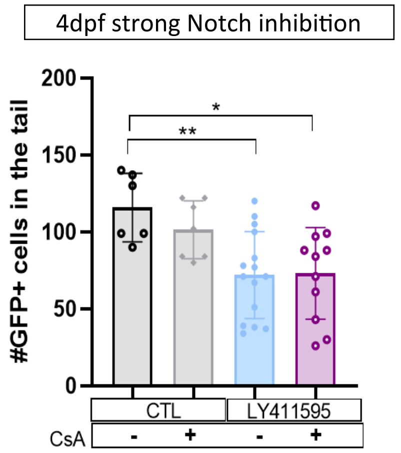

### Supplemental figure 6

##### Figure 6 supplemental

A

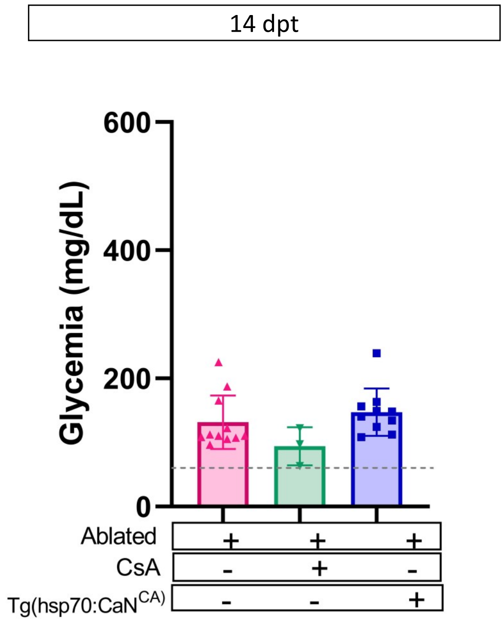


B

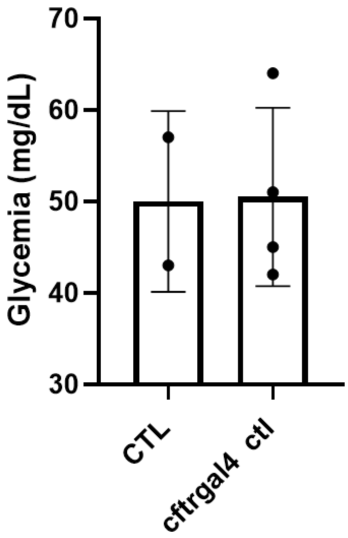
